# Network analysis of correlated conformational dynamics across the evolution of influenza A virus NS1

**DOI:** 10.64898/2026.09.29.755463

**Authors:** Abhishek Bastiray, James E. Gonzales, Wonmuk Hwang, Jae-Hyun Cho

## Abstract

Protein evolution requires sequence changes to alter molecular properties without disrupting the dynamic organization necessary for function. How residue-level networks of correlated motions accommodate such mutations remains poorly understood. Here, we investigate the evolution of correlated conformational dynamics in nonstructural protein 1 (NS1) variants of influenza A viruses that emerged between 1918 and 2004. From molecular dynamics (MD) simulations we built residue networks in which edge weights reflect generalized correlations of conformational motion. The network of correlated dynamics is characterized using centrality analysis. Overall network properties are well-conserved across NS1 variants, indicating evolutionary robustness of the dynamical network. Residues ranked highly across multiple centrality measures form a predominantly hydrophobic cluster in the protein core. These residues exhibited low conformational entropy and high sequence conservation, consistent with rigid, coordinated motions. In contrast, residues exhibiting large differences in betweenness and closeness centrality ranks occupied flexible, solvent-exposed regions surrounding the core network. These residues also showed lower evolutionary conservation. These findings reveal a core-periphery structure in which a rigid hydrophobic network enables long-range dynamic connectivity while more adaptable peripheral regions accommodate evolutionary sequence variation. This architecture provides a mechanistic framework for long-range epistatic interactions without compromising the underlying structural framework.

## Introduction

Evolution encodes patterns in protein sequences and structures.^1-3^ However, how these patterns emerge and change during evolution remains incompletely understood. Thus, uncovering the principles underlying the evolutionary patterns is important for understanding protein evolution at the molecular level.

Mutations drive protein evolution by constantly reshaping interactions between residues.^4^ A protein does not populate a single static structure but instead exist as a dynamic ensemble of conformations.^5, 6^ Characterizing the dynamics underlying these conformational ensembles is essential for understanding the regulation of protein functions, including protein-protein interactions, allostery and post-translational modifications.^7-9^ To fully understand how mutations reshape protein function during evolution, mechanisms for mutational effects on protein dynamics need to be understood. Despite recent advances in machine learning-based approaches,^10, 11^ quantitative prediction of mutational effects on protein dynamics remains challenging.

Although short-range mutational effects can often be rationalized through structural comparisons between wild-type and mutant proteins, the mechanisms underlying long-range effects on protein dynamics are much more difficult to identify.^12-15^ A useful first step in studying long-range mutational effects is to compare the correlated dynamics between all residue pairs in wild type and mutant proteins.^16, 17^ However, pairwise correlations alone often provide limited information about how correlated residues are directly or indirectly connected across a protein structure. Network analysis provides a useful framework for revealing the connectivity underlying these correlated residues.^18-20^

There are several approaches for constructing the intramolecular protein communication network. For example, networks can be constructed based on residue-residue contacts in a single static structure or across structural ensembles.^21^ Alternatively, networks can be based on inter-residue correlations of conformational dynamics.^13, 22^ Residue coevolution can also be used to construct intramolecular networks.^23, 24^ These approaches capture distinct but potentially complementary aspects of intramolecular interactions.

In the present study, we investigated how evolutionary mutations affect the network of intrinsic conformational dynamics across a protein structure. Our network analysis is based on generalized correlation of conformational dynamics (GCCD) between residues, calculated from molecular dynamics (MD) simulation trajectories.^25^ In this study, nodes correspond to residues, whereas edges represent the strengths of generalized correlations in conformational dynamics between residue pairs.

Centrality is a measure of the topological importance of nodes or edges within the overall network structure.^26^ Although many centrality measures have been developed to characterize distinct network properties, four measures are commonly applied in the analysis of biological networks, betweenness, closeness, degree, and eigenvector centralities. Brief descriptions of these centralities are provided below. In the network used in this study, shorter path lengths, or distances, correspond to stronger correlations of conformational dynamics between residues.

Betweenness centrality (BC) measures how often a residue lies on the shortest paths between all other pairs of nodes in a network.^27^ A residue with high BC often acts as a key node through which many shortest paths pass as they traverse the network. Closeness centrality (CC) measures how close a residue is to all other residues in a network, based on its shortest path lengths to them.^28^ A residue with high CC has short network distances to all other residues, so such a residue participates in global conformational dynamics across the protein. Degree centrality (DC) measures the number of edges connected to a residue.^28^ A residue with high DC is directly connected to many other residues and shares correlated motions with many immediate network neighbors. Eigenvector centrality (EC) measures the importance of a residue based not only on its number of direct connections, as for DC, but also on the centralities of other residues connected to it.^29^ A residue with high EC is connected to other residues that are highly correlated with others. Such a residue could serve as a hub for allosteric communication.

Because each centrality measure reflects a distinct aspect of network topology, combining multiple centrality measures can provide a more comprehensive view of network architecture. In the present study, we compare the four centrality measures to: (1) characterize the overall network structure of correlated dynamics, (2) determine how evolutionary mutations affect the network structure, and (3) identify residues that have distinct roles in connecting correlated dynamics across the protein structure. Together with structural analyses, this network-based approach provides insight into the differential roles of network core and peripheral residues in long-range correlated dynamics.

Nonstructural protein 1 (NS1) of influenza A virus (IAV) is a major viral virulence factor.^30^ NS1 binds to a number of host proteins to antagonize innate immune responses.^31^ Structurally, NS1 consists of an RNA-binding domain (RBD) and an effector domain (ED), followed by a disordered C-terminal tail (**Fig 1A**).^32^ Although full-length NS1 forms a homodimer mediated by the RBD, the isolated ED forms a monomer in solution and retains its function in binding host proteins.^12, 33^ Consequently, the isolated NS1 ED has served as a useful model for investigating the molecular mechanisms underlying NS1-mediated viral immune evasion.^34^

**Figure 1.**
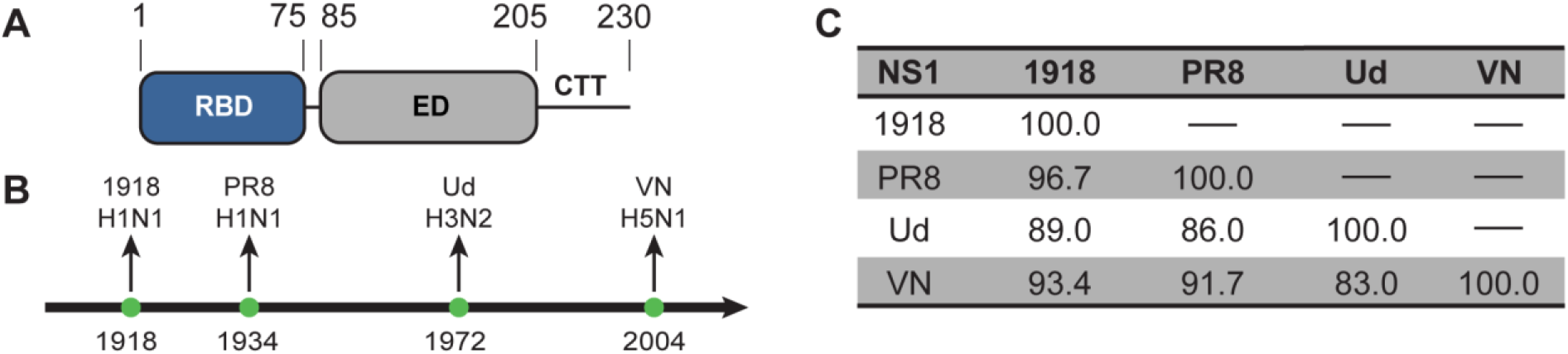
Nonstructural protein 1 (NS1). (**A**) Domain organization of NS1. (**B**) Timeline of the emergence of the four IAV variants. (**C**) Sequence variations among the four NS1 variants.

NS1 is also one of the most frequently mutated proteins in the IAV genome.^35^ In the present study, we employed four NS1 ED variants (hereinafter NS1) from IAV strains emerged between 1918 and 2004: 1918 H1N1, Puerto Rico 8 (PR8) H1N1, Udorn (Ud) H3N2, and Vietnam (VN) H5N1 IAVs (**Fig 1B**). These NS1 variants contain 3 – 17 % of sequence variations among them (**Fig 1C**).^12^ Despite these sequence variations, major biological functions of these NS1 variants are conserved. Therefore, comparison of these variants provides an opportunity to investigate how evolutionarily near-neutral mutations reshape the biophysical properties and dynamical organization of NS1 while preserving its overall structural and functional framework.

Our previous study identified long-range correlated dynamics involving hydrophobic core residues in 1918 and PR8 NS1 varaints.^12, 13^ However, the PR8 IAV, which emerged in 1934, is an immediate descendant of 1918 IAV, and their NS1s differ by only four mutations.^13, 36^ Therefore, how long-range correlated dynamics evolved over a longer timescale remains unclear.

In this study, we examine the robustness of the network roles of hydrophobic core residues, particularly their role in long-range correlated dynamics during NS1 evolution. Overall, our study provides mechanistic insight into how conformational dynamics are coupled among residues across the NS1 structure.

## Results

### Structural features of high centrality residues

We calculated four centrality measures for NS1 variants, 1918, PR8, Ud, and VN, derived from IAVs that emerged between 1918 and 2004. Residues were ranked in descending order of centrality score, with rank 1 representing the highest-ranked, most influential residue. We then identified the top 10-ranked residues for each centrality measure.

We noted that a subset of residues was ranked highly across all four centrality measures. Specifically, although the top 10 residues for the four centrality measures could theoretically comprise up to 40 distinct residues in each NS1 variant, on average, only approximately 20 unique residues were identified (**Fig 2A**). Because these residues were repeatedly identified across different centrality measures, they occupy important positions in the network. We therefore termed these Group A residues and examined their structural and chemical features (**Fig 2B**).

**Figure 2.**
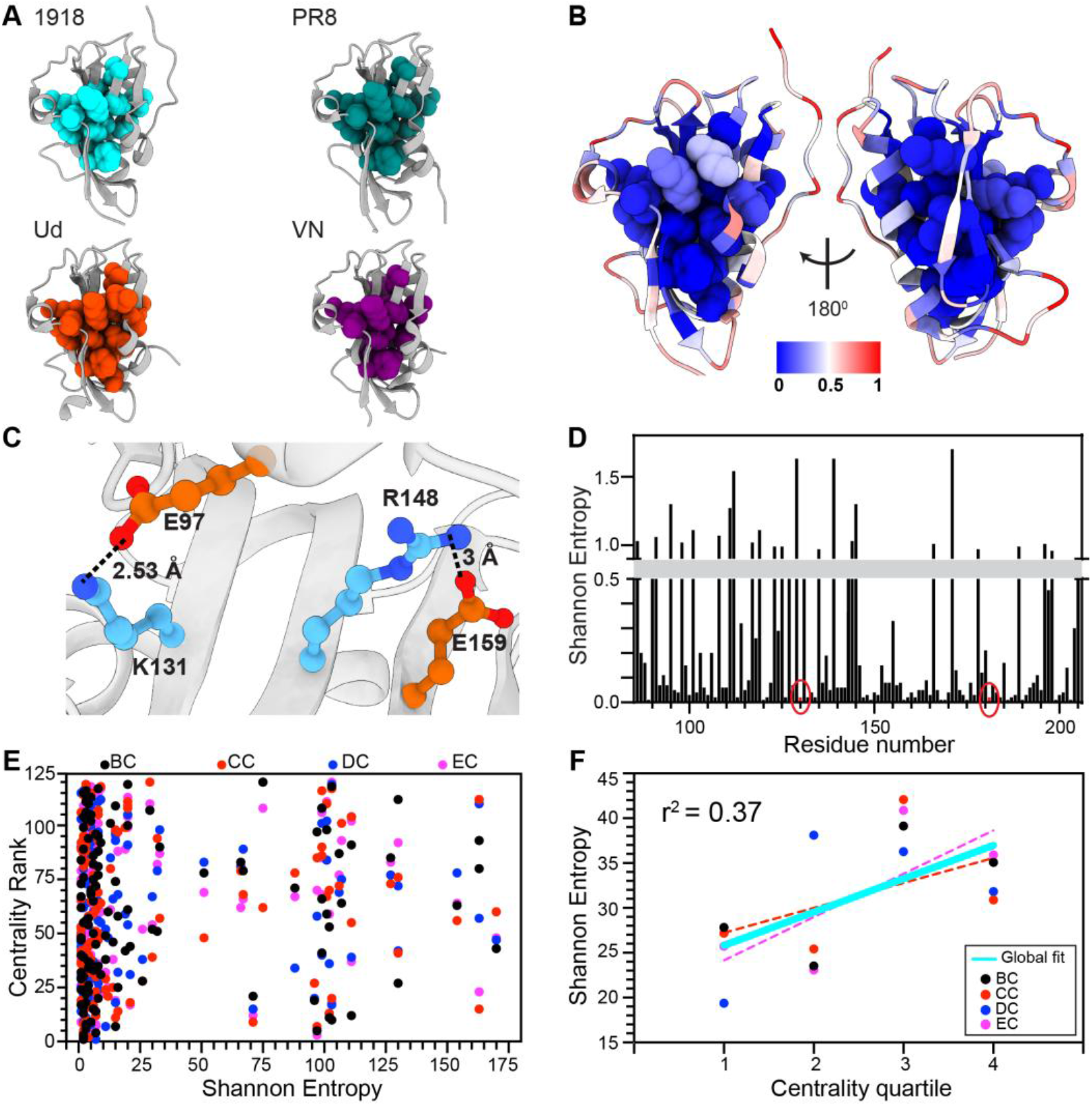
Central residues in the NS1 network. (**A**) Residues ranked in the top 10 by four centrality measures. (**B**) Positions of Group A residues in the NS1 structure. (**C**) Salt-bridges formed by Group A residues. (**D**) Shannon entropy as a function of residue numbers. Residues 130 and 181 are circled in red. The gray rectangle indicates the break in the y-axis. (**E**) Centrality rank as a function of Shannon entropy of individual residues in 1918 NS1. The corresponding results for the other NS1 variants are shown in Supplementary Figure 1 (Fig S1). (**F**) Shannon entropy as a function of the centrality quartile of individual residues in 1918 NS1. The corresponding results for the other NS1 variants are shown in Fig S2.

Group A consisted predominantly of hydrophobic residues and lacked polar residues. Group A residues were largely buried in the NS1 structure, with an average relative solvent accessible surface area (rSASA) of 5.3 ± 3.8 %, compared with 48 ± 1 % for all other residues (**Fig 2B**). This result suggests that close physical packing contributes to strong correlations of inter-residue conformational dynamics, consistent with our previous results^12, 13^ and those of others.^37^ Interestingly, three Group A residues across the four NS1 variants were charged amino acids, residues 131, 148, and 159, all of which are partially solvent exposed. Specifically, R148 forms a salt bridge with E159, and K131 forms a salt bridge with E97 which is highly conserved in NS1 (**Fig 2C**). Although charged residues have usually been associated with low centrality values,^38^ these charged Group A residues may occupy high-centrality positions because of their structural and functional importance.^39-42^

Notably, two Group A residues, L130 and I181, appeared among the top-ranked residues for all four centrality measures in all NS1 variants. Thus, these two residues are likely to be the most critical nodes in the network of dynamic correlations. Their potential importance is further supported by their near-complete conservation, as indicated by low Shannon entropy, among all nonredundant human IAV NS1 sequences collected between 1918 and 2020 (n = 5,574) (**Fig 2D**). Despite the high conservation of Group A residues, there was no apparent general correlation between centrality scores and Shannon entropy because many low-centrality residues are also highly conserved (**Fig 2E and Fig S1**). To examine overall relationship between centrality and residue conservation, we assigned residues to quartiles according to their centrality ranks, with the first quartile for the top 25% ranked residues. We then calculated the average Shannon entropy of residues within each quartile.

The relationship between centrality-rank quartiles and average Shannon entropy revealed a weak but significant trend in which higher-centrality residues were more highly conserved (**Fig 2F and Fig S2**). The same pattern was observed across all four centrality measures and NS1 variants. This finding is consistent with previous reports of an inverse relationship between residue centrality and evolution rate.^38^ Moreover, it was reported that highly conserved residues are significantly more spatially clustered than randomly selected residues.^43, 44^ Nevertheless, the relatively weak correlation observed here suggests that centrality is only one of multiple factors influencing residue conservation. Other factors, including requirements for ligand or protein-binding specificity, may also contribute substantially to mutation rates.^45^

### Group A residues are conformationally rigid

Group A residues form a cluster in the hydrophobic core of NS1 (**Fig 2B**), suggesting that their conformational dynamics is restricted by tight packing. To quantify their flexibility, we calculated residue-specific conformational entropy for both backbone and side chains. To eliminate the dependence of side-chain entropy on the number of rotatable bonds, side-chain entropy was normalized by the number of side-chain dihedral angles before being combined with backbone entropy.

Group A residues exhibited significantly lower average conformational entropy than the rest of residues, with 8.59 ± 0.11 and 9.34 ± 0.02 kcal/mol at 300K for Group A and the remaining residues, respectively. Centrality rank was also associated with conformational entropy, such that more conformationally rigid residues tended to have higher centrality ranks (**Fig 3A and Fig S3**). The same pattern was observed in all four centrality measures across the NS1 variants.

**Figure 3.**
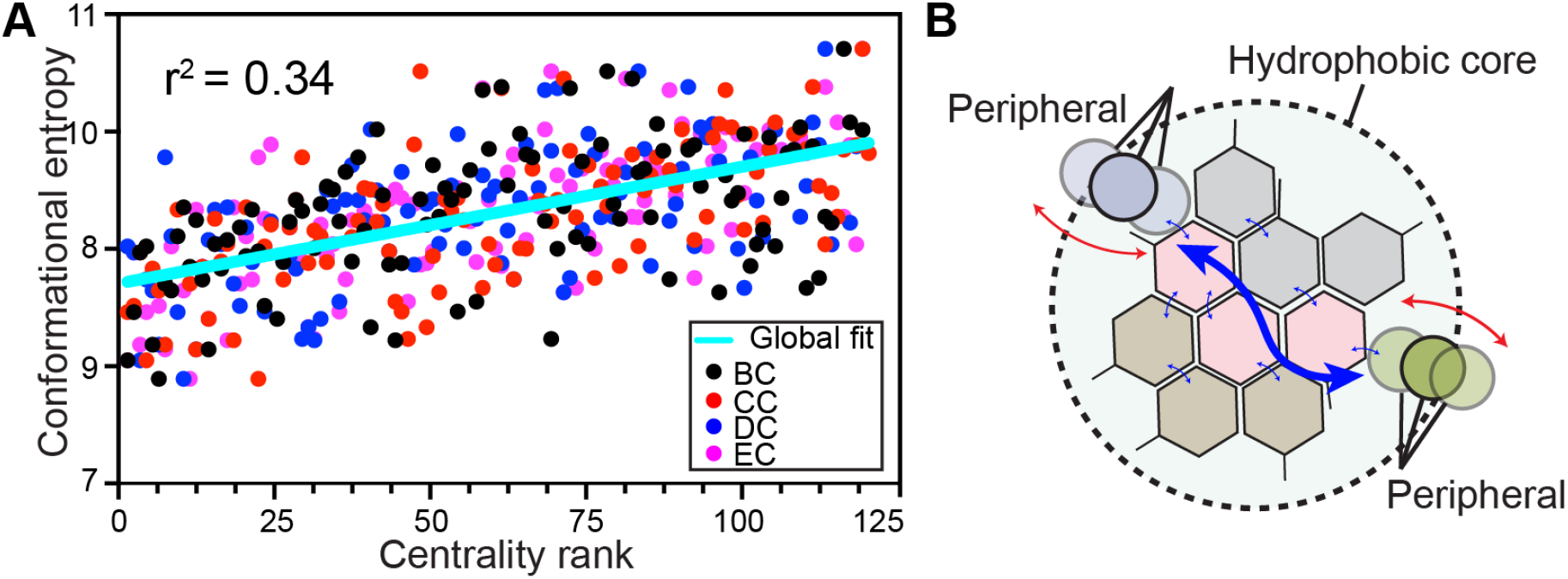
Characteristics of Group A residues. (**A**) The contribution of conformational entropy to the free energy at 300K (kcal/mol) for individual residues in 1918 NS1 as a function of centrality rank. The corresponding results for the other NS1 variants are shown in Fig S3. (**B**) Schematic showing how concerted dynamics of Group A residues contribute to long-range correlated dynamics. The blue arrow represents the epistatic interaction between two distant peripheral residues.

These observations suggest that the central region of the dynamical network consists mainly of conformationally rigid residues. Dense packing restricts the conformational space accessible to these residues while maintaining close physical interactions through which their residual fluctuations can remain correlated with those of neighboring residues. Consequently, rigid residues such as Group A can form a stable, densely interconnected network characterized by relatively small but coordinated conformational fluctuations (**Fig 3B**). Concerted motions of hydrophobic core residues have also been observed in other proteins.^46-50^

However, these results should not be interpreted as indicating that all Group A residues constitute a single dynamically coherent block. Group A was defined by combining residues selected across multiple centrality measures and NS1 variants (Fig 2A). Therefore, different subsets of Group A residues may form distinct but overlapping dynamical clusters, producing a mosaic-like structure of correlated motions.

The correlated dynamics involving Group A residues may also contribute to long-range epistasis, in which the effect of one mutation depends nonlinearly on the presence of another mutation at a distant site. In our previous NMR study, mutations at distant sites in NS1 affected one another through small but detectable changes in the structure and conformational dynamics of hydrophobic core residues. These observations are consistent with the present network analysis and support a potential role for the hydrophobic core in coupling the effects of distant mutations across the NS1 structure.

### Peripheral network regions frequently harbor evolutionary mutations

Correlations among different centrality measures provide insight into overall network architecture. Moreover, comparing these relationships across protein variants can reveal how network architecture is conserved or altered during evolution. Therefore, we examined the correlations between centrality measures across four NS1 variants. Overall, the centrality measures for each NS1 variant correlated well with each other (**Figs 4A and 4B**). The mean correlation coefficient across all pairs of centralities and the four NS1 variants was 0.8. The consistently high correlations across the four variants suggest that overall relationships among different centrality features are evolutionarily robust.

**Figure 4.**
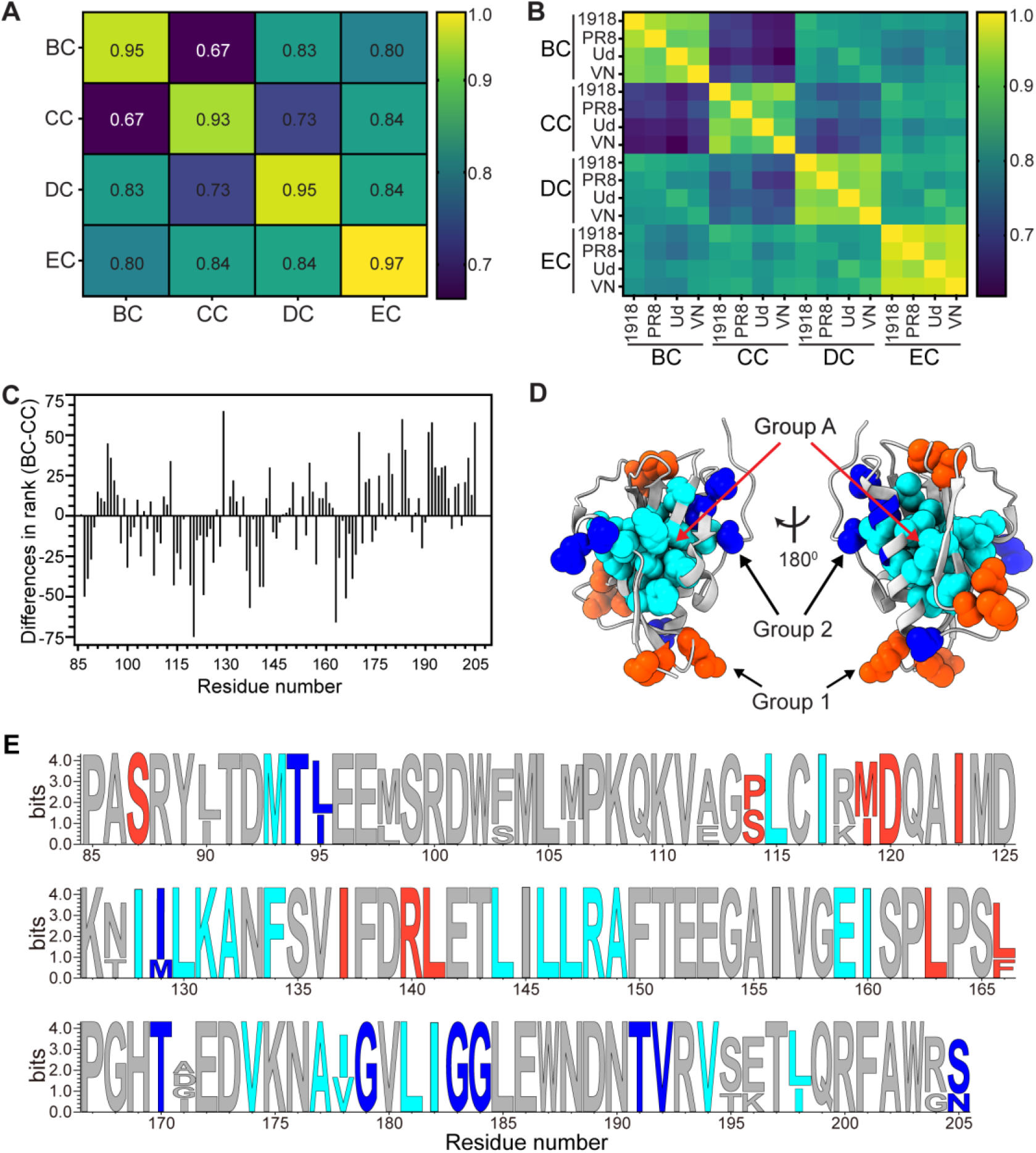
Relationships among centrality measures. (**A**) Heatmap of the average correlation coefficients between centrality measures. (**B**) A 16 × 16 heatmap of the correlation coefficients between centrality measures across four NS1 variants. (**C**) Δr as a function of residue numbers of 1918 NS1. The Δr of other NS1s are in Fig S4. (**D**) Positions of Group A (cyan), Group 1 (orange) and Group 2 (blue) residues in the 1918 NS1 structure. The positions of these groups in the other NS1 structures are shown in Fig S5. (**E**) Sequence logo showing Group A, Group 1 and Group 2 residues for 1918 NS1. The corresponding results for the other NS1 variants are shown in Fig S6.

However, BC and CC showed the least correlation among all pairs of centralities across the four NS1 variants, with an average correlation coefficient of 0.67 (**Fig 4A**). This relatively weak correlation reflects substantial differences between BC and CC ranks for a subset of residues.

These residues may play distinct roles in the network from those of the central core residues, which rank highly in both BC and CC. We therefore examined these residues and their underlying structural and functional features.

For each NS1 variant, we first identified the 10 residues with the largest negative and the 10 residues with the largest positive differences between their BC and CC ranks (**Fig 4C and Fig S4**).

Δr = rBC - rCC

where rBC and rCC denote residue ranks based on BC and CC centrality scores, respectively. Residues were ranked in descending order of centrality score, such that rank 1 corresponds to the highest centrality score. Consequently, a smaller rX (X = BC or CC) indicates a higher rank for centrality measure X. Residues with large negative differences (Δr < 0) have higher BC ranks than their CC ranks and were designated Group 1. Conversely, residues with large positive differences (Δr > 0) have higher CC ranks than their BC ranks and were designated Group 2.

Notably, neither Group 1 nor Group 2 residues were in the top 10 ranks in either BC or CC, except residues 119, 129, and 194. These three residues were ranked either 9th or 10th in BC or CC for the PR8, Ud, or VN NS1 variants (**Figs 4D-E and Figs S5-S6**). Rather, they generally occupied intermediate-to-low ranks in BC or CC. Specifically, the average BC and CC ranks for Group 1 were 39 and 91, respectively, whereas those of Group 2 were 90 and 37, respectively, across four NS1 variants.

These results indicate that Group 1 residues participate in relatively more shortest paths than would be expected from their relatively large average network distances to other residues. Namely, Group 1 residues exhibit greater betweenness relative to their global network proximity. In contrast, Group 2 residues have relatively short average network distances to the rest of residues but lie on relatively fewer shortest paths between other residues. Thus, they are comparatively well integrated into the network but do not serve as major intermediaries along shortest paths.

These contrasting features suggest distinct roles of Groups 1 and 2 in the network. The combination of relatively high BC and low CC for Group 1 is consistent with a bridge-like role connecting peripheral regions to the core network. Mutations at these positions could therefore preferentially perturb correlations between specific network regions without strongly disrupting the central network. In contrast, the relatively low BC of Group 2 suggests that these residues are less important for shortest-path routing across the network. Mutations at Group 2 positions may have more localized effects within the global network of correlated dynamics. Based on these properties, we hypothesized that Groups 1 and 2 may be more tolerant to evolutionary mutations than Group A.

To test this idea, we examined sequence variability at these positions during NS1 evolution. The average Shannon entropy values for Group 1 and 2 across all human IAVs were 0.37 and 0.39, respectively, significantly higher than that for Group A residues (0.24). We next examined structural features of Groups 1 and 2 that might contribute to their higher mutation frequency than Group A. Indeed, Groups 1 and 2 are located in distinctive positions from Group A residues in the NS1 structure (**Fig 4D**). Specifically, Groups 1 and 2 are located primarily in flexible, solvent-exposed loop regions of NS1 and form multiple smaller clusters spatially separated from one another. Noticeably, these clusters all lie in peripheral regions immediately outside the central Group A cluster. In contrast, Group A residues form a single cluster in the hydrophobic core and are embedded in rigid regions of the protein structure.

Moreover, the four selected NS1 variants differ by 3% - 14% in sequence in pairwise comparisons (**Fig 1C**). These mutated sites largely overlap with Groups 1 and 2. Specifically, the mean sequence distance from each mutation site to the nearest Group 1 or Group 2 residue was only 0.9 residue positions (**Fig 4E** and **Fig S6**). These observations suggest that evolutionary mutations are concentrated near the network periphery while avoiding the highly conserved core. This finding is also consistent with the observed correlation between residue centrality and the mutation rate (**Fig 2E**).

Previous studies have also reported higher mutation rates at topologically peripheral residues than at core residues.^38, 51-54^ However, it should be noted that network core and periphery can be defined using different criteria, and no single definition is universally applicable to inter-residue interaction networks. In the present study, we did not assign residues to core or peripheral regions a priori. Rather, this organization emerged from the combined analysis of multiple centrality measures together with their locations within the structure. Future studies will be required to determine whether this approach consistently identifies structurally and topologically meaningful core–periphery organization in other protein residue networks.

## Discussion

In this study, we investigated the residue-level network of correlated conformational dynamics across four NS1 variants from IAVs that emerged between 1918 and 2004. Despite sequence variations among them, the overall network architecture was largely conserved, suggesting that the organization of correlated dynamics are evolutionarily robust.

The core and peripheral network residues exhibited markedly different structural and network properties. The core residues (Group A) form a single cluster within the hydrophobic core of NS1. In contrast, Groups 1 and 2 form multiple peripheral clusters that are spatially separated from one another and located around the core network. Group A residues are conformationally rigid due to dense packing of hydrophobic side chains, whereas Groups 1 and 2 are generally more solvent-exposed and flexible. Moreover, Group A residues are highly conserved, while Groups 1 and 2 residues exhibit higher mutation rate during IAV evolution.

These findings suggest that the tightly packed hydrophobic core may provide a structural scaffold that supports long-range dynamic coupling between distant residues. Mutations among the four NS1 variants occurred predominantly at or near, in sequence, residues belonging to the peripheral groups, which also overlap protein–protein interaction surfaces. Perturbations introduced by mutation or protein binding at these peripheral sites may therefore be dynamically coupled to distant regions through the rigid core of the network. This organization provides a potential mechanistic framework for epistasis.^12, 13, 55^ Mutations with small individual effects may reshape the network of correlated motions and alter the functional effects of mutations at other sites.^13^

Previous studies showed that long-range interdomain communication in multidomain proteins can be mediated by relatively sparse networks of residues, including key residues located at the interdomain or hinge positions.^20, 42, 56, 57^ In contrast, the present study examined how correlated motions are organized within a single-domain architecture. Our findings therefore complement hinge models by describing how conformational dynamics are connected within domains on either side of hinge positions.

An important caveat of the present study is the timescale of the protein dynamics examined. Our analysis primarily characterizes fast, sub-microsecond fluctuations. These motions predominantly reflect fluctuations within local conformational substates and do not directly describe slower, barrier-crossing transitions between distinct conformational basins.

Nevertheless, fast backbone and side-chain motions contribute to conformational entropy and can influence protein thermodynamics, including stability and binding free energy^58, 59^. Future studies integrating longer-timescale dynamics with mutational and functional analyses will be needed to determine how the conserved NS1 network connects fast correlated motions to evolutionary changes in protein function.

## Methods

### MD simulations

We prepared five independent systems for the MD simulations for the 1918 (PDB ID: 6U28^33^, resolution: 2.95 Å), PR8 (PDB ID: 3O9S^60^, resolution: 2.48 Å), Ud (PDB ID: 3KWG^61^, resolution: 2.21 Å), and VN (PDB ID: 3P31^62^, resolution: 2.45 Å) NS1 systems. Protocols for setting up the solvated system and performing simulations follow the same method as described previously.^13^ Briefly, CHARMM^63-65^ (version c48a1 with the param36 all-atom force field^66^) was used to build, solvate, and neutralize in a cubic water box of TIP3P water.^67^ Sodium (Na^+^) and Chloride (Cl^-^) ions were added to neutralize the system at a concentration of 50 mM. A 6-stage energy minimization procedure was then used on the systems.^13^ The systems were then heated to and equilibrated at 300 K before performing production MD simulations. The production simulations were carried out using OpenMM version 7.6.0 for 1 μs.^68^

### Contact Network Analysis

The contact network analysis follows the same protocol as previously described.^13^ Briefly, the network utilized the generalized dynamic cross-correlation coefficients to compute the edge lengths between nodes in the graph.^20^ Edges were included in the graph if the residues formed a high-occupancy contact (> 70%) during the trajectory. The NetworkX package in Python^69^ and Mathematica version 14.3 (Wolfram) were used to perform the graph theory analysis.

### Mutation analysis of the NS1 amino acid sequence

All human H1N1 IAV sequences (5,574 non-duplicated sequences) were obtained from the NCBI influenza research database (https://www.fludb.org/brc/home.spg?decorator=influenza). The sequences were aligned using the MUSCLE (Multiple Sequence Comparison by Log-Expectation) algorithm.^70^

## Acknowledgements

Research reported in this publication was supported by the National Institute of General Medical Sciences of the National Institutes of Health under grant number R35GM152007 (to J.H.C.) and by the Welch Foundation under grant number A-2028-20260402 (to J.H.C.). A total of

$1,832,739, or 84% of the project costs, was financed with federal funds. A total of $351,000, or 16% of the project costs, was financed with non-governmental funds. The content is solely the responsibility of the authors and does not necessarily represent the official views of the National Institutes of Health.

