## Supplementary Information for "Network analysis of correlated conformational dynamics across the evolution of influenza A virus NS1"

**Figure S1**

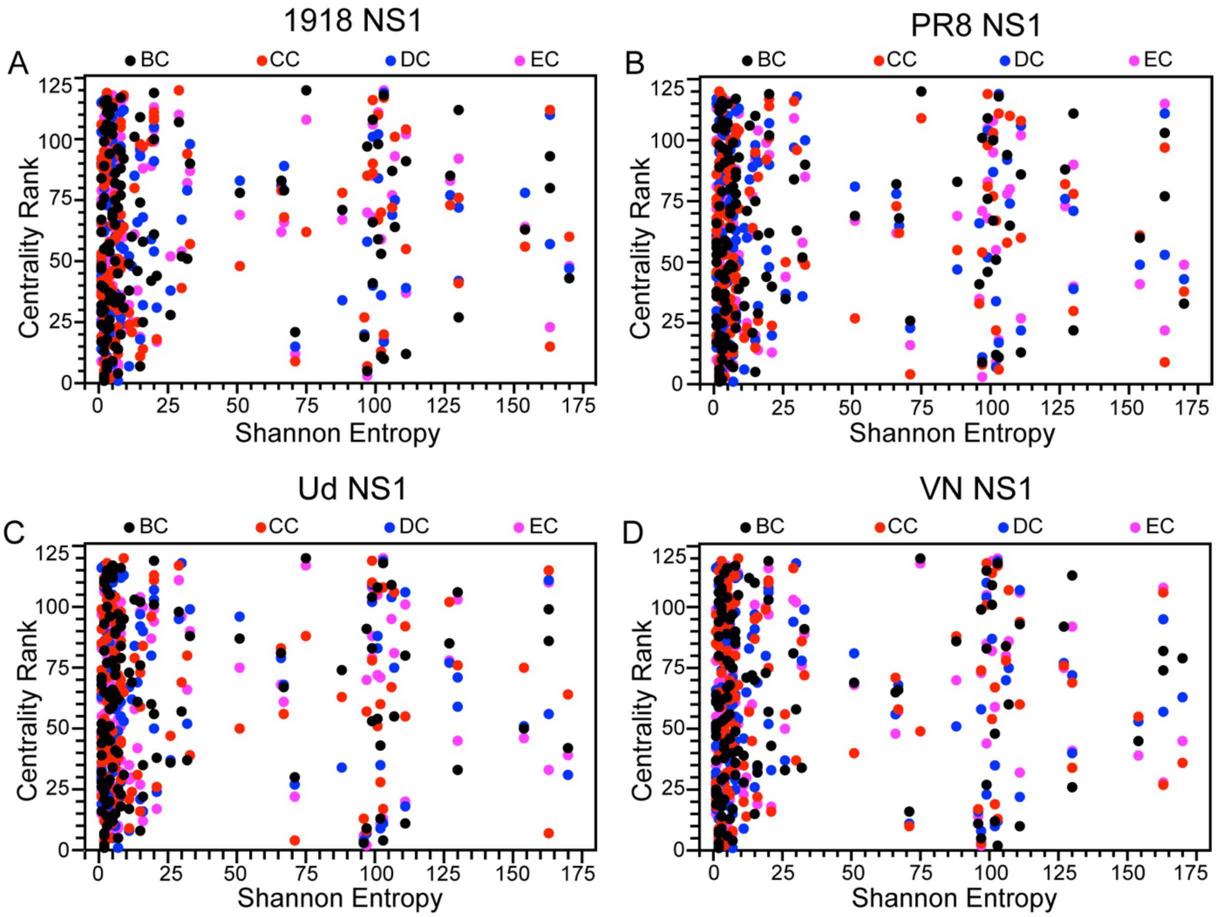


**Supplementary Figure 1.** Centrality rank of residues of individual NS1 as a function of Shannon entropy **(A)** 1918 NS1 **(B)** PR8 NS1 **(C)** Ud NS1 **(D)** VN NS1.

**Figure S2**

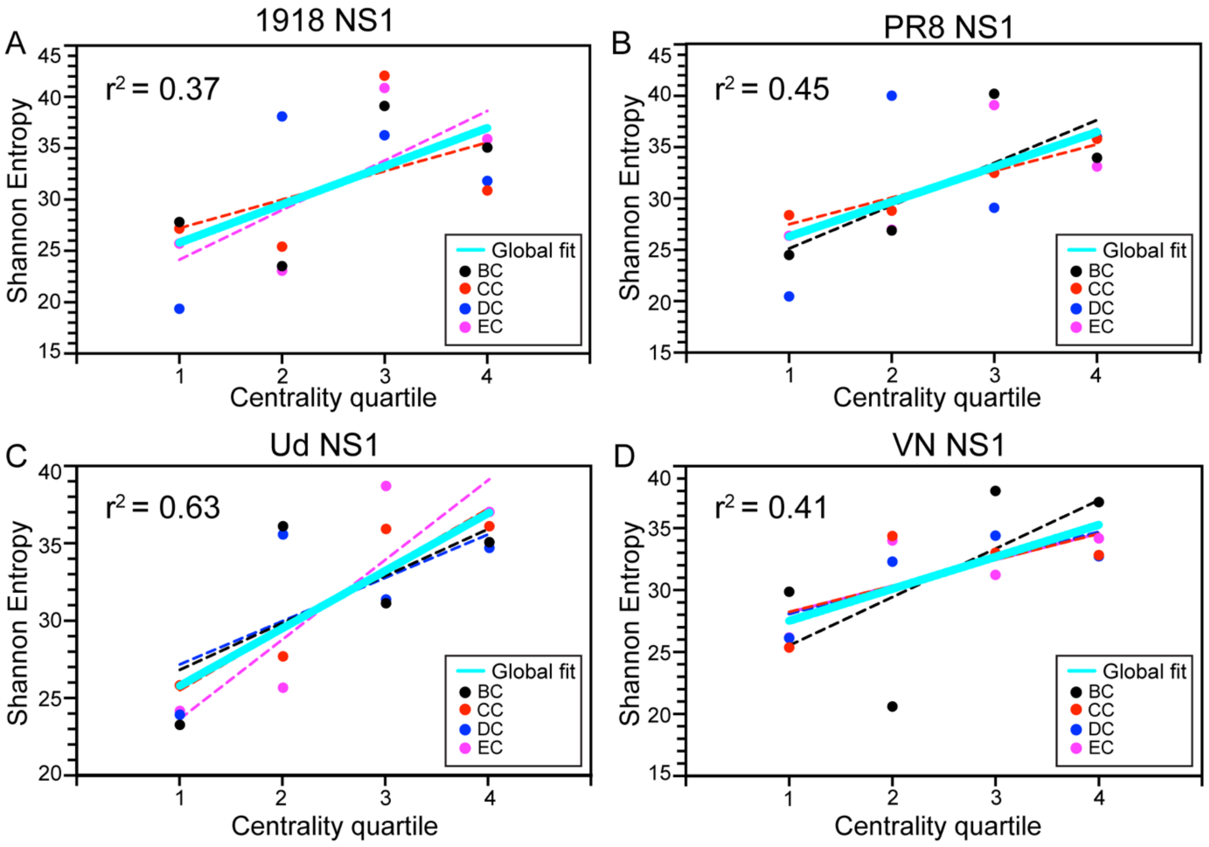


**Supplementary Figure 2.** Shannon entropy of residues of individual NS1 as a function of centrality quartile **(A)** 1918 NS1 **(B)** PR8 NS1 **(C)** Ud NS1 **(D)** VN NS1. Dashed lines represent linear fits for the individual centrality measures, whereas solid line represents the global fit across all centrality measures. The $r^2$ value of each global fit for mentioned in the top left corner of each panel.

**Figure S3**

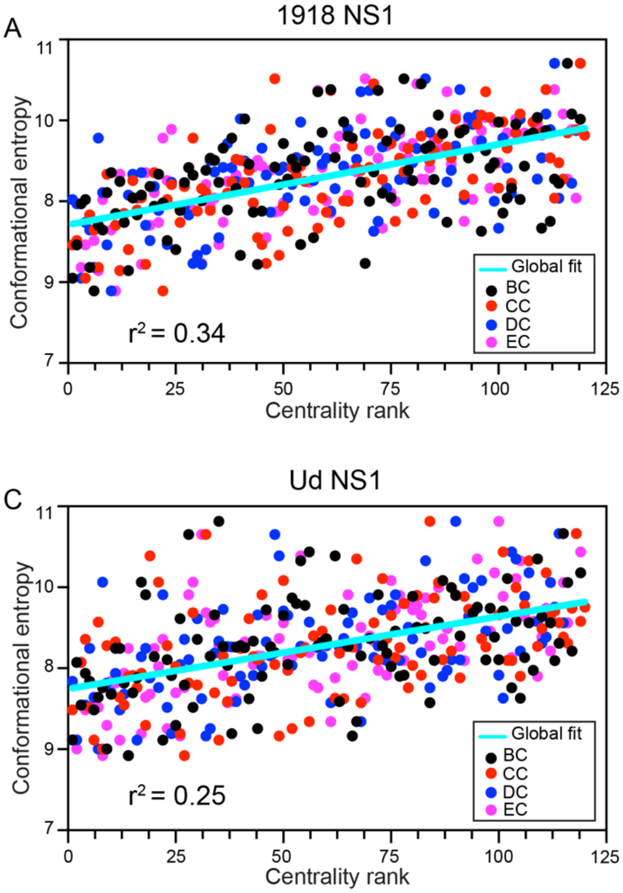


**Supplementary Figure 3.** Conformational entropy of Group A residues of individual NS1s as a function of centrality rank **(A)** 1918 NS1 **(B)** PR8 NS1 **(C)** Ud NS1 **(D)** VN NS1. The solid line represents the global linear fit across all centrality measures, while corresponding $r^2$ value of global fit is mentioned in the bottom left corner of each figure.

**Figure S4**

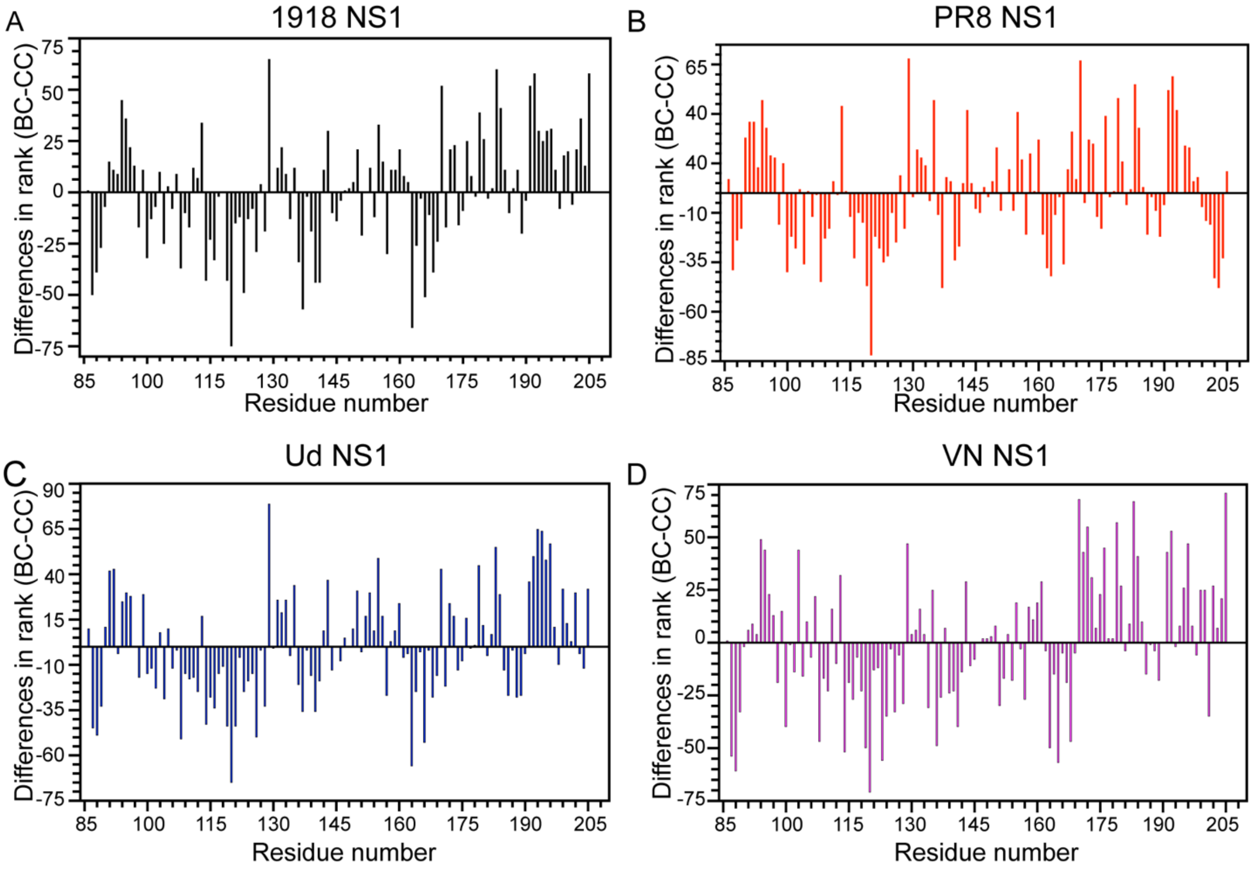


**Supplementary Figure 4.** Difference in between betweenness and closeness centralities rank (BC-CC rank **;(**Δr)) as a function of residue numbers for different NS1s **(A)** 1918 NS1 **(B)** PR8 NS1 **(C)** Ud NS1 **(D)** VN NS1.

**Figure S5**

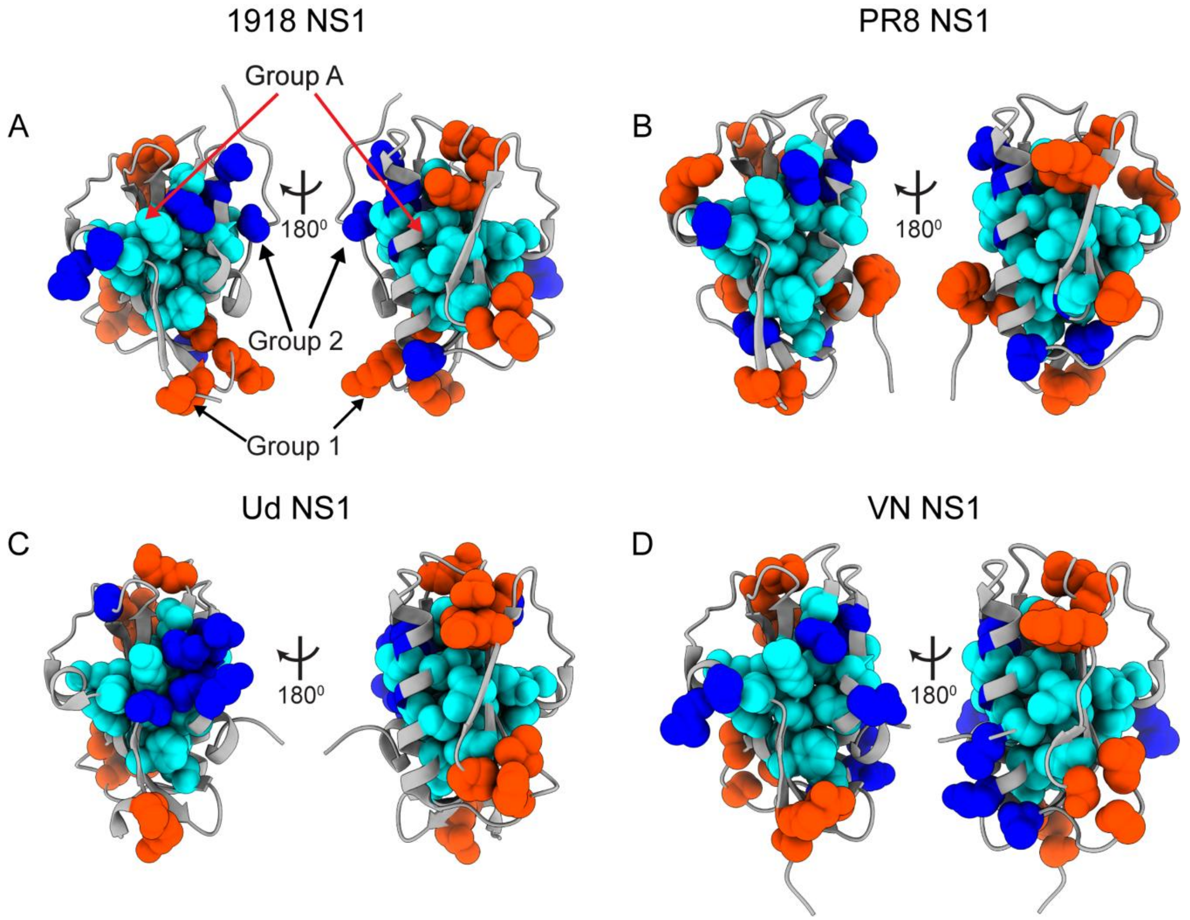


**Supplementary Figure 5.** Structural distribution of Group A (cyan), Group 1 (orange red) and Group 2 (blue) residues in **(A)** 1918 NS1 **(B)** PR8 NS1 **(C)** Ud NS1 **(D**) VN NS1. Two orientations related by $180^0$ rotation are shown for each NS1 structure.

**Figure S6**

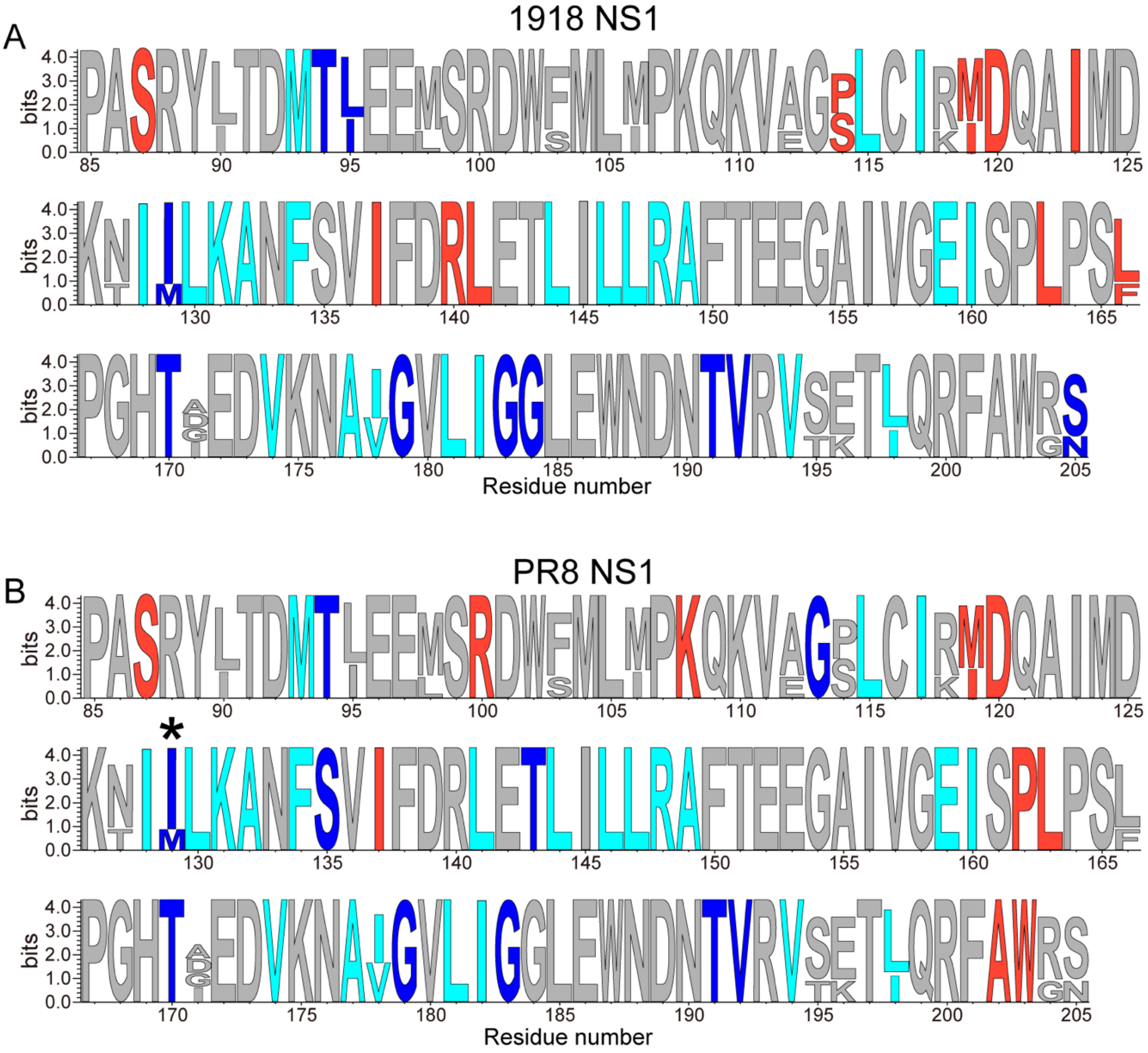

**Figure S6 (continued)**

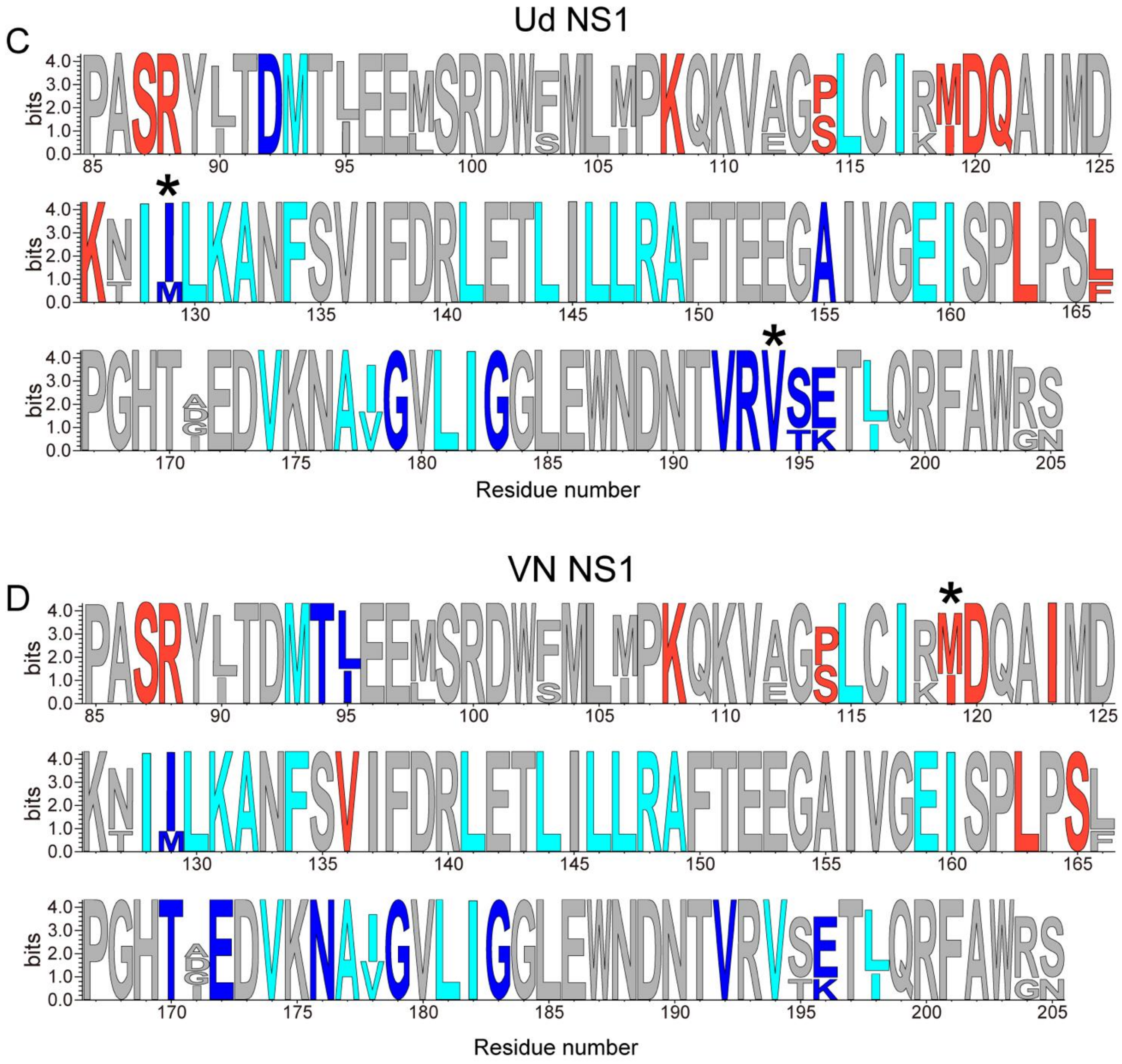


**Supplementary Figure 6.** Sequence logo of NS1 showing Group A (in cyan), Group 1 (red orange) and Group 2 (in blue) residues for **(A)** 1918 NS1 **(B)** PR8 NS1 **(C)** Ud NS1 **(D)** VN NS1. Asterisks (*) indicate residues that belong to Group A and either Group 1 or 2.
